# Micro-RNA 7975 directly regulates MDTH expression and mediates endothelial cell proliferation and migration in the development of early atherosclerosis

**DOI:** 10.1101/2024.10.15.618502

**Authors:** Nathan Wagner, Genesio Karere

## Abstract

Cardiovascular disease (CVD) is commonly due to the development of atherosclerosis. Endothelial integrity is critical in the prevention of pathogenesis of atherosclerosis. The key to prevention of CVD is understanding the molecular mechanisms responsible for initiation of early atherosclerosis. MiRNAs are mediators of endothelial homeostasis, and their dysregulation could lead to early atherosclerotic disorder. We previously revealed the expression of miR-7975 in early atherosclerotic lesions. The aim of this study was to investigate the novel roles of miR-7975 on endothelial cell proliferation and migration, and in the regulation of metadherin (MTDH) expression. We performed proliferation and migration assays coupled with luciferase assay. We show that miR-7975 promotes proliferation and migration of endothelial cells and that miR-7976 directly regulates (MTDH), previously associated with cancer pathogenesis. In conclusion our results show miR-7975 could be a potential mediator of endothelial homeostasis and that MTDH is a novel target of miR-7975.

## INTRODUCTION

Atherosclerotic cardiovascular disease is the leading cause of deaths in the world and accounts for nearly half of the deaths (Benjamin et al. 2018) and produces immense health and economic burdens (Martin et al. 2024). Atherosclerosis is progressive disease accompanied by multifaced processes initially characterized by lipoprotein oxidation (LO), endothelial activation, injury, and dysfunction, allowing infiltration of LO and monocytes into the subendothelial layer of the vascular tissue (Libby P 2011). Hallmarks of endothelial pathophysiological processes involved in early atherosclerosis, include proliferation and migration disorder, apoptosis, reactive oxygen production (ROS) and inflammation (Albarran-Juarez et al. 2016; Li et al. 2021; Wu et al. 2022; Cheng et al. 2023). Thus, the maintenance of endothelial homeostasis is critical in the prevention of initiation of early-stage atherosclerosis, and lack of, could ultimately results in acute ischemic events (Choy et al. 2001). However, the underlying mechanisms that drive initiation and progression of early-stage atherosclerosis remain unclear. Elucidating the molecular mechanisms that underlie development of early-stage atherosclerosis is critical for identification of potential therapeutic targets for early intervention prior to development of life-threatening unstable plaques.

One key mediator of atherosclerosis is a microRNA (miRNA), a small noncoding RNA that regulate gene expression post-transcriptionally, fine-tunning multiple cellular processes involved in atherogenesis, including endothelial cell proliferation and migration. miRNAs perform their functions by binding to 3’untranslated regions of massager RNA (mRNA) resulting in either degradation of mRNA or inhibition of translation. The resultant effect of miRNA functions is decrease of protein abundance in the cell. Numerous studies have implicated miRNA mediated mechanisms in the endothelial cell processes in atherosclerosis pathway (e.g., (Zhang et al. 2020; Li *et al*. 2021). Previously we reported the development of early atherosclerosis in baboons (N=112) changed with high cholesterol, high fat (HCHF) diet for two years (Mahaney et al. 2018). After evaluating the burden of atherosclerosis in formalin-fixed common iliac arteries, we used small RNA sequencing to identify all expressed miRNAs in a paired fashion: early-stage lesions (fatty streaks and fibrous plaques) and adjacent health regions (control) of the same tissue, providing a self-control study design. We reveled several differentially expressed miRNAs in lesions relative to control, including miR-7975 which significantly upregulated in both fatty streak and plaques lesions (Karere et al. 2023). In parallel, we identified predicted miRNA gene targets, including metadherin (MTDH targeted by miR-7975 (Karere *et al*. 2023). While miR-7975 and MTDH are implicated in cancer development (Zhou et al. 2012; Liu & Yang 2013; Du et al. 2014; Howe et al. 2022; Mimmi et al. 2023; Wang et al. 2024), to our knowledge, neither miR-7975 nor MTDH are associated with the development of atherosclerosis. Hence, the elucidation of the regulatory role of miR-7975-MTDH axis to mediate atherosclerosis is novel.

In the present study, we explored the role of miR-7975 in proliferation and migration of endothelial cells. Our data revealed that miR-7975 directly regulates MTDH and inhibit cell proliferation and migration. Our study provides novel findings of potential therapeutic targets for effective treatment of early atherosclerosis.

## METHODS

### Cell culture, treatment and transfection

Primary human umbilical endothelial cells (HUVEC; Lonza Catalog #: C2519A) were cultured in Lonza EBM-2 media (CC-3155) with the Lonza EGM-2 supplement kit (CC-4176). Cells were be cultured in a humified atmosphere with 5% CO_2_ at 37oC until 60 to 80% confluency.

### Luciferase assay

To evaluate whether miR-7975 directly regulates MTDH expression, we performed luciferase assay. Luciferase reporter vector was synthesized by Azenta Inc. Briefly, segments of the MDTH’s 3’UTR-miRNA target sites or a mutant target sequence were subclone into a pGL3 vector with SV40 promoter for transient expression of luciferase gene (Promega). We performed targeted sequencing to validate the insert and construct. We used pRL-TK Renilla vector with thymidine kinase promoter (Promega) as a control vector. For luciferase assay, we used Lipofectamine 3000 (Thermal Fisher) to co-transfect HepG2 cells with miR-7975 oligonucleotide mimics (60nM) and Luciferase vector with insert (wild type, wt,) or Renilla control vector in two groups: Luciferase-wt + miR-7975 mimic + Renilla (assay) and Luciferase-wt + untargeting miRNA mimic + Renilla (negative control). Cells were cultured for 24 h before performing luciferase assay using Dual-Glo Luciferase/Renilla Assay System (Promega) and Infinite 200 Pro plate reader (Tecan Ltd) to determine transfection efficiency and luciferase activity relative to the control. Data was normalized against Renilla activity. All transfections were carried out in triplicates.

### Proliferation assay

HUVEC were grown in T75cm^2^ flasks, trypsinized and replated into a 96 well plate and allowed to grow to 70% confluency. Once at the desired confluency, the cells were transfected with a miR-7975 mimic using Lipofectamine 3000 following the manufacturer’s protocol. Transfection media was replaced with complete growth media after 6 hours. The cells were then allowed to incubate for up to 72 hours post transfection. At 24, 48, and 72 hours post transfection XTT proliferation assays were performed according to the manufacturer’s protocol. Briefly, an XTT working solution was made and then immediately added to the cells’ growth media. After 4 hours of incubation, absorbance at 450nm was measured for actively proliferating cells and at 660nm for background signal from cell debris. The final formula for calculating the specific absorbance is: [Abs450 nm (Test) – Abs450 nm (Blank)] – Abs660 nm (Test). Absorbance (Test) is from wells with cells, and Absorbance (Blank) is from wells that contain only media and the XTT reagent.

### Migration assay

HUVEC cells were grown in T75cm^2^ flasks, trypsinized and replated onto gelatine coated glass coverslips in a 12 well plate and allowed to grow to 70% confluency. Once at the desired confluency, the cells were transfected with a miR-7975 mimic using Lipofectamine 3000 following the manufacturer’s protocol. Immediately after transfection, a p200 pipette tip was used to scratch a line through the center of the cells growing on the glass coverslips. Images of the cells were taken at 0, 24,48, and 72 hours post transfection. ImageJ was used for image processing and analysis.

### Statistical analysis

Statistical analysis was performed using two tailed student t-test or ANOVA followed by Tukey’s post hoc test, p < 0.05.

## RESULTS

### Luciferase assay

miR-7975 directly regulate MTDH expression as demonstrated by the results of the luciferase assay. The luciferase activity in HepG2 cells cotransfected with miR-7975 mimic and a vector carrying segment of the miRNA binding site on MTDH 3’UTR (assay) was significantly different relative to control (p=0.03) and decreased by approximately by 40% (Figure 1).

**Figure 1:**
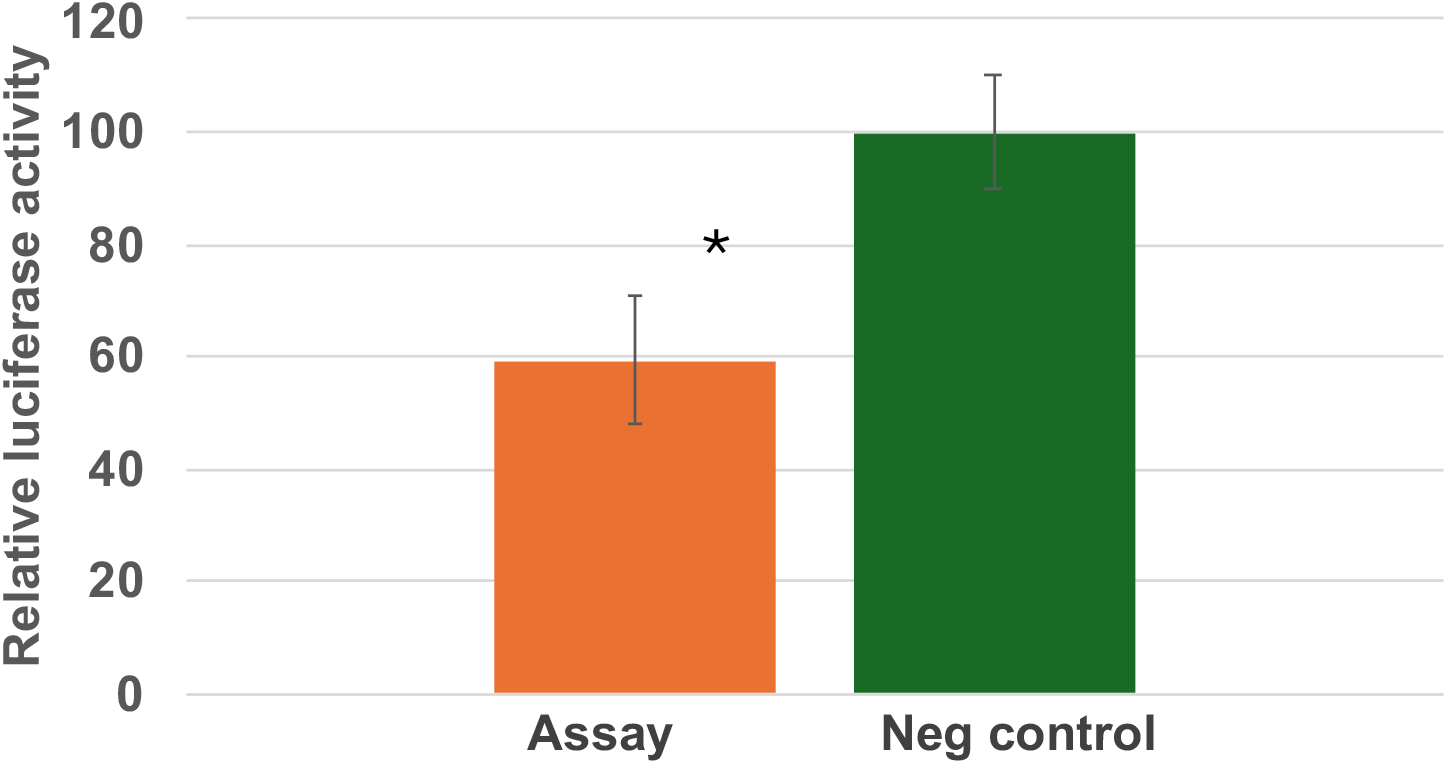
Experimental validation of miR-7975 interaction with MTDH. pGL3 vector construct with 3’UTR insert was transfected into HepG2 in two experiments: Assay was the vector construct containing 3’UTR and miR-7975 mimic and the negative control was the construct and untargeting miRNA mimic. Renilla vector (pRL-TK) was used as an internal control. Results are shown as relative luciferase activity, with the control defined as 100%. Bars show mean±SEM of the experiments, each performed in triplicate. ^*^ Indicates p<0.05.

### Proliferation assay

miR-7975 promoted HUVEC proliferation as shown by the results of the XTT assay over time. Proliferation of the cells was not significantly different between the treated cells and control at 24 hrs (p=0.43) post transfection of miR-7975. However, the difference was profound at 48 (p=0.01) and 72 hrs (p=0.02) post transfection (Figure 2).

**Figure 2:**
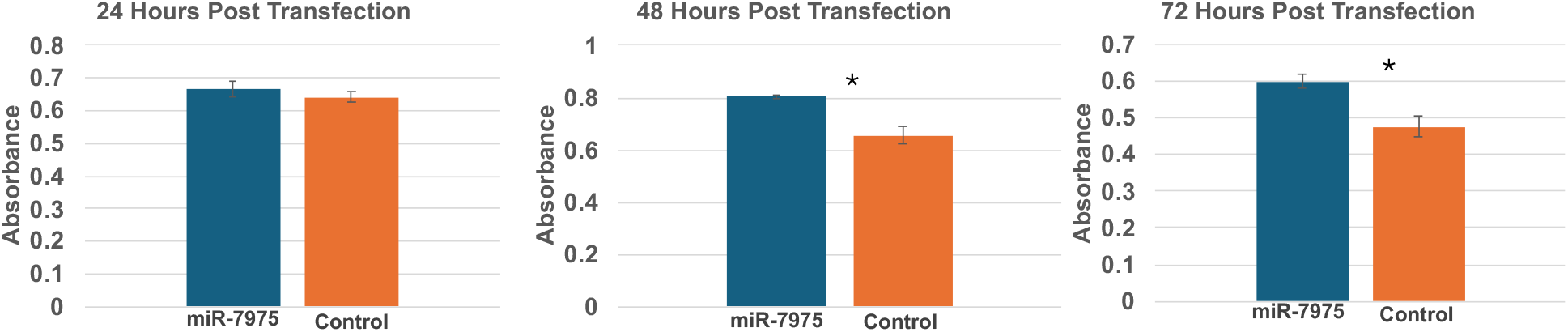
miR-7975 promoted HUVEC cell proliferation. Proliferation was evaluated by performing XTT assay at 24, 48, and 72hrs post transfection with miR-7975 mimic compared to control. ^*^ Indicates p<0.05.

### Migration assay

miR-7975 promoted HUVEC migration as shown by the results of the scratch assay over time. Figure 3 demonstrates the migration of the cells towards the scratch area at 24, 48, and 72hrs post miR-7975 mimic transfection compared to control. The results show a decreasing trend of the scratch area over time. Figure 4 is a graphical presentation of the recovered scratch area over time post transfection.

**Figure 3:**
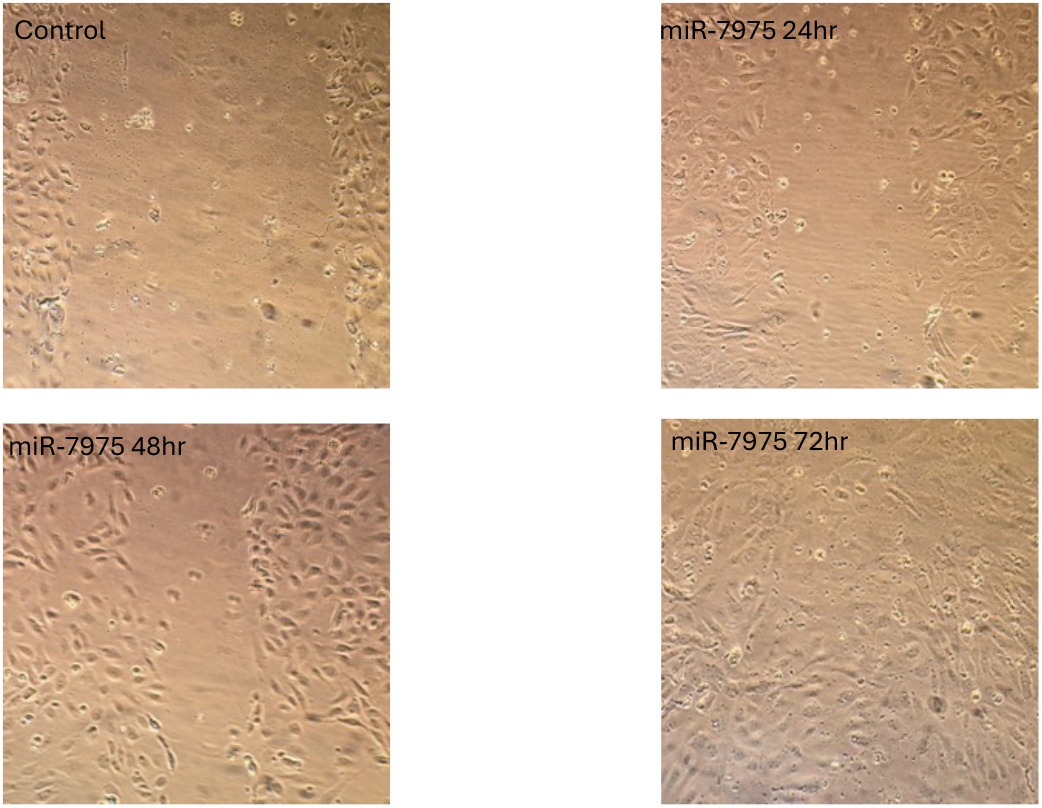
miR-7975 enhanced HUVEC migration compared to control. We performed the migration assay and evaluated the scratch area at 24, 48, and 72hrs post transfection of miR-7975 mimic. The results show a decreasing trend of the scratch area over time.

**Figure 4:**
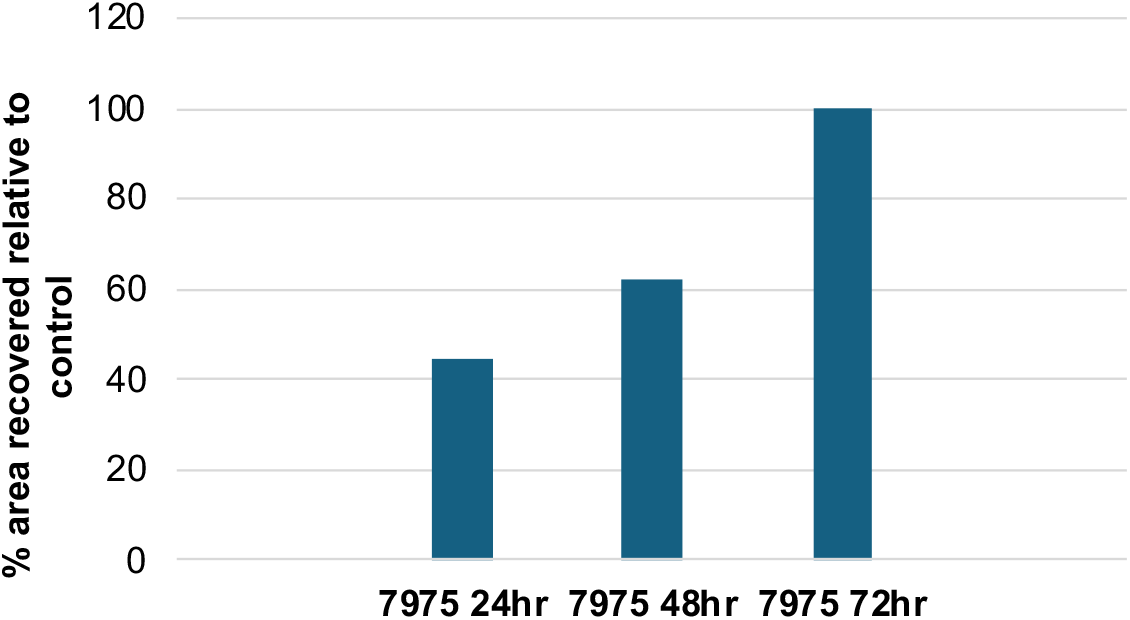
Graphic representation of % recovery of scratched area during HUVEC migration. We performed the migration assay and evaluated the scratch area at 24, 48, and 72hrs post transfection of miR-7975 mimic. The results show a decreasing trend of the scratch area over time.

## DISCUSSION

Cardiovascular disease (CVD), a leading cause of deaths globally, is commonly due to the development of atherosclerosis (Benjamin *et al*. 2018) and produces immense health and economic burdens (Martin *et al*. 2024). Endothelial integrity is critical in the prevention of pathogenesis of atherosclerosis (Limbu & McCloskey 2024). The key to prevention of CVD is understanding the molecular mechanisms that drive initiation of early atherosclerotic lesions prior to their progression to life-threatening plaques. Our results show miR-7975 could be a potential mediator of endothelial homeostasis and that MTDH is a novel target of miR-7975.

In our previous study we challenged baboons with HCHF diet for 2 years and evaluated the extent and early atherosclerotic lesion types (fatty streaks or fibrous plaques) (Mahaney *et al*. 2018). We then investigated the molecular changes associated with development of lesions in common iliac arteries of baboons (Karere *et al*. 2023). We revealed that miR-7975 expression was significantly upregulated between paired lesions and adjacent health regions of same tissues. These findings led us to the obvious corollary that the miR-7975 may have a potential functional role in development of early atherosclerotic lesions.

Few studies have investigated miR-7975 roles in disease development but to our knowledge, no studies have associated miR-7975 with the development of atherosclerosis. A study investigating the molecular changes associated with abuse of 3,4-Methylenedioxy methamphetamine (MDMA) revealed that the expression of miR-7975 is suppressed in ventral tegmental region of the brain for MDMA abuser compared to control, suggesting that the miRNA may play a role in response to use of MDMA (Demirel *et al*. 2019). In contrast, miR-7975 was upregulated in cerebrospinal fluid of patients with lung adenocarcinoma and that the miRNA promoted cell proliferation, migration and invasion, indicating that the miRNA may be potential biomarker and therapeutic target against lung adenocarcinoma (Pan *et a*l. 2020). Other studies have shown that miR-7975 is contained in neutrophil extracellular vesicles, and is proinflammatory (Youn *et al*. 2021), negatively correlated with gestational age at birth (Howe *et al*. 2022), and upregulated in serum exosomes positive for angiotensin-converting enzyme 2 (ACE2) in patients with COVID19 (Mimmi *et al*. 2023). Although the miR-7975 has not been associated with atherosclerosis, the findings of the previous studies suggest that it is involved in regulation of cellular processes that are hallmark of atherosclerosis: proinflammatory, promotion of cell migration and proliferation. Our findings confirm miR-7975 regulates these cellular processes in vascular endothelial cells, a major cell type involved in maintaining vascular integrity (Choy *et al*. 2001).

In the previous study, we used bioinformatic miRNA target prediction tools, including TargetScan, TarBase and miRecords implemented in Ingenuity Pathway Analysis (IPA) and identified MTDH as potential target of miR-7975 (Karere *et al*. 2023). MTDH, localized in human chromosome 8, codes for metadherin protein. A few studies that investigated the roles of the protein have shown that metadherin is an oncogene involved in the development of multiple cancer types, including gallbladder adenocarcinoma (Liu & Yang 2013), hepatocellular carcinoma (Wang *et al*. 2024) and bladder cancer (Zhou *et al*. 2012). Other cancer biology studies have indicated that the protein mediates trastuzumab regimen resistance in breast cancer patients by decreasing PTEN abundance via -NFkB pathway (Du *et al*. 2014). To our knowledge, no studies have investigated the role of MTDH in the development of atherosclerosis, suggesting it is a novel therapeutic target for effective treatment of the disease.

## CONCLUSIONS AND FUTURE STUDIES

In our study, we investigated the mechanistic role of miRNA-MTDH pathway in the development of early atherosclerosis. We have shown miR-7975 directly targets MTDH and that miR-7975 promotes vascular endothelial cell migration and proliferation. Future studies will investigate the roles of MDTH in vascular endothelial and smooth muscle cell processes related to atherosclerosis as well as in macrophages using silencing and overexpression approaches in atherosclerotic cell models, and subsequently in preclinical animal models.

## ACKNOWLEDGMENTS

This work was supported by the National Institutes of Health (K01HL130697) received by GMK and the Wake Forest University Claude D. Pepper Older Americans Independence Center (P30-AG021332).

